# LESSeq: Local event-based analysis of alternative splicing using RNA-Seq data

**DOI:** 10.1101/841494

**Authors:** Jing Leng, Christopher JF Cameron, Sunghee Oh, James P Noonan, Mark B Gerstein

**Affiliations:** Program in Computational Biology and Bioinformatics, Yale University, New Haven, CT, USA; Department of Molecular Biophysics and Biochemistry, Yale University, New Haven, CT, USA; Division of Human Genetics, Department of Pediatrics, Cincinnati Children’s Hospital Medical Center, Cincinnati, OH, USA; Department of Genetics, Yale University School of Medicine, New Haven, CT, USA; Kavli Institute for Neuroscience, Yale University School of Medicine, New Haven, CT, USA; Department of Computer Science, Yale University, New Haven, CT, USA

**Keywords:** RNA-Seq, alternative splicing, Maximum Likelihood Estimation

## Abstract

Alternative splicing, which can be observed genome-wide by RNA-Seq, is important in cellular development and evolution. Comparative RNA-Seq experiments between different cellular conditions allow alternative splicing signatures to be detected. However, inferring alternative splicing signatures from short-read technology is unreliable and still presents many challenges before biologically significant signatures may be identified. To enable the robust discovery of differential alternative splicing, we developed the Local Event-based analysis of alternative Splicing using RNA-Seq (LESSeq) pipeline. LESSeq utilizes information of local splicing events (i.e., the partial structures in genes where transcript-splicing patterns diverge) to identify unambiguous alternative splicing. In addition, LESSeq quantifies the abundance of these alternative events using Maximum Likelihood Estimation (MLE) and provides their significance between different cellular conditions. The utility of LESSeq is demonstrated through two case studies relevant to human variation and evolution. Using an RNA-Seq data set of lymphoblastoid cell lines in two human populations, we examined within-species variation and discovered population-differential alternative splicing events. With an RNA-Seq data set of several tissues in human and rhesus macaque, we studied cross-species variation and identified lineage-differential alternative splicing events. LESSeq is implemented in C++ and R, and made publicly available on GitHub at: https://github.com/gersteinlab/LESSeq

## Introduction

Alternative splicing of precursor-messenger RNA (pre-mRNA) generates multiple transcripts (or isoforms from a single gene locus) that may differ in localization, function or other biological features. Alternative isoform usage is thought to be a major source of biological complexity during development and evolution (1). In humans, alternative splicing variations have been implicated in differential disease associations and drug responses (2). These implications highlight the need for a deeper understanding of the associations, or even causal relationships, between alternative splicing and human biological variations. During evolution, alternative splicing leads to transcriptome (and sometimes proteome) expansion in organisms through differential inclusion and exclusion of exonic sequences. This expansion is believed to underlie lineage-specific phenotypic traits (1).

Over the past few years, high-throughput RNA sequencing or RNA-Seq (3) has dramatically expanded our knowledge of alternative splicing. Almost all human multi-exonic pre-mRNAs were discovered to undergo alternative splicing and that tissue-specific regulation of alternative splicing may be pervasive (4), which suggests a functional relevance for alternative isoform usage. However, it has also been shown that noisy products from alternative splicing are extensive (5), emphasizing the need to distinguish biologically important alternative splicing events from those of no functional consequences, which could be achieved in part through comparative RNA-Seq experiments that include multiple biological conditions.

Computationally, many methods have been developed to assemble and quantify RNA transcripts from RNA-Seq data. Since the transcriptome of any given condition is unlikely to be fully captured by a reference annotation, it is desirable to assemble condition-specific transcripts by leveraging available RNA-Seq data. Expression levels of the newly assembled transcripts can then be calculated before downstream analysis (e.g., differential transcript usage detection between two conditions). However, there are many challenges that exist when inferring transcripts from RNA-seq data. First, assembling the correct isoforms from short-read data for the sample(s) of a study is very difficult, especially for mammalian genomes (6). In most transcript assembly methods designed for mammalian genomes, RNA-Seq reads are first mapped to the reference genome. The resulting exonic and spliced reads are then used (sometimes in conjunction with a reference transcriptome annotation) to construct a splicing graph for each gene locus. This gene locus is then used to derive isoform structures according to a specific graph-traversal algorithm. Since short-read technology does not provide full connectivity of different regions in the splicing graph, the strategy for traversing such graphs varies wildly across methods (i.e., generate the most parsimonious set of graphs, all possible graphs, or somewhere between these two extremes), and none have been shown to yield satisfactory full-transcript annotation in human (6). Second, even if the ‘correct’ transcriptome annotation is provided, determining the level of expression for each transcript is not trivial (7) as short-read technology necessitates probabilistic estimation of transcript abundances. Most transcript quantification tools calculate the Maximum Likelihood Estimate (MLE) of transcript abundances based on a specific objective function, whose form and complexity differ across methods and are dependent on the modeling of RNA-Seq data. In addition, transcript quantification algorithms based on similar ideas can generate different results (6), with agreement between algorithms generally decreasing as the number of isoforms of a gene increases (8). Moreover, the assembly and quantification steps are tightly linked, as incorrect transcript annotation exacerbates the quantification problem (8). Combined, the inaccuracies and uncertainties in both transcript assembly and quantification make transcript-based comparative study extremely challenging.

We thus propose an upstream approach to the transcript-based RNA-Seq inference problem. Here, we devise a Local Event-based analysis of alternative Splicing using RNA-Seq (LESSeq) analytical approach that focuses on nearby regions in genes, where isoform structures may diverge (e.g., skipped or included exon encompassed by two constitutive exons) (4, 9). By assessing local alternative splicing events, our method sidesteps several aspects of uncertainty in transcript-based analysis and generates more robust results. Local alternative splicing events are essentially local parts of splicing graphs that contain diverging paths, and analyzing such regions abrogates the need to accurately reconstruct full-length transcripts. Since transcript assembly methods output different transcripts even with the same underlying splicing graph, circumventing the step of whole transcript assembly bypasses the errors produced from it. In addition, the number of local events in a given gene is never greater than that of all the isoforms of a gene, thus yielding more robust quantification results - as has been shown previously, fewer isoforms per gene lead to more consistent quantification results between methods (8). Moreover, focusing on a set of defined, simple patterns for local events yields regions that are guaranteed to be computationally identifiable (10), which is not always true for transcript-level inference.

## Materials and methods

The four major steps of the LESSeq pipeline are described below (see Fig. 1A).

**Fig. 1.**
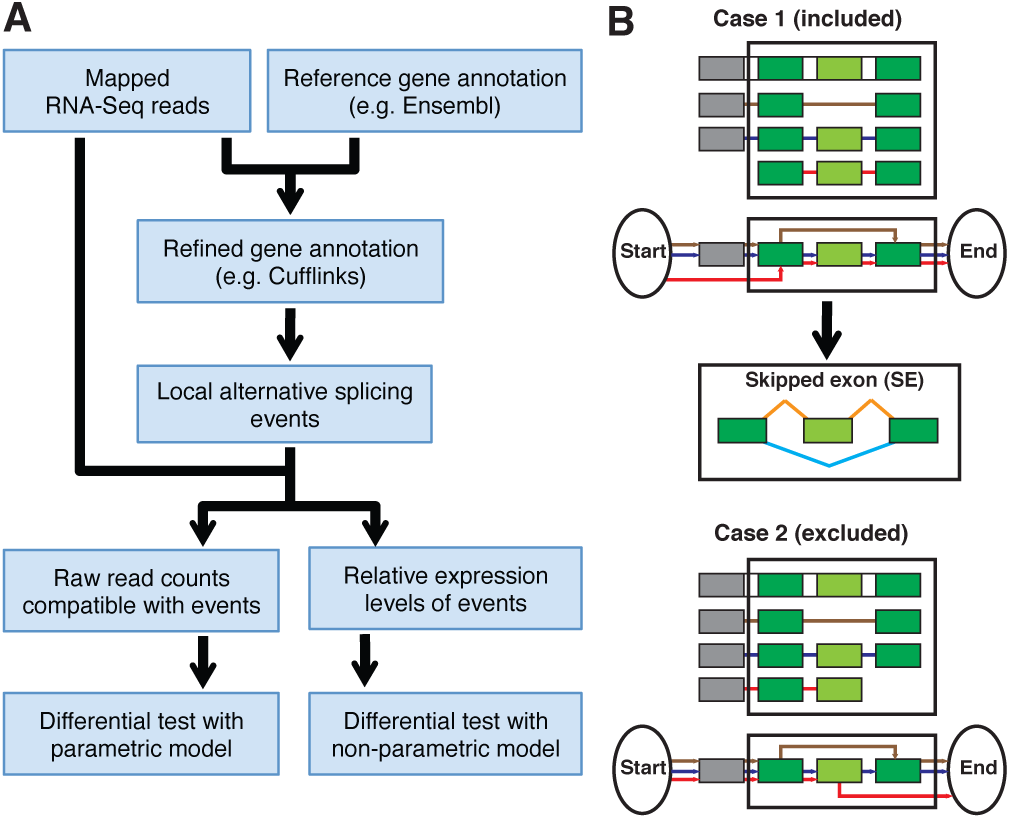
Local event-based analysis of RNA-Seq data. (**A**) Pipeline overview. (**B**) Local event identification. A splicing graph is built for each gene locus, with each isoform represented as one path in the graph (red, blue and brown lines – panel **B**). In LESSeq, a local event (in black lined boxes) is identified such that the out most two exons (dark green) are present in all isoforms (i.e., constitutive), with all in- and out-edges going through the 5’ and 3’ exons, respectively. The shortest such local graphs are always taken, so that no other constitutive exons are present within the identified local graphs. Such a conservative definition of local events ensures the accurate quantification of each alternative form of the local events. In the example shown in Case 1, three isoforms collapse into two alternative forms of a local ‘skipped exon’ event. Constitutive and variables exons are colored in dark and light green, respectively. Orange and cyan lines represent the connection between exons in two alternative forms of a local event. Case 2 represents a local event (3’ constitutive exon missing) that is excluded by the LESSeq pipeline.

### Refining gene models using RNA-Seq data

LESSeq’s first step is to derive comprehensive splicing annotations for the specific sample(s) of a study. For species with a reference annotation, this step is optional but strongly recommended when deep sequencing data is available. Alternative splicing is highly tissue-specific and there may exist splicing events in the condition of interest that are not annotated in a reference transcriptome. The current implementation of LESSeq employs Cufflinks (11) for splicing annotation, where the Reference Annotation Based Transcript assembly (RABT option) for well annotated organisms (e.g., human) is recommended. With the RABT method, faux-reads that tile the reference transcripts are used together with RNA-Seq reads to assemble splicing structures. However, alternatives to Cufflinks may also be used. For species that do not have a reference genome and/or reference transcriptome annotation, de novo methods to build transcript models are available (12). Users should substitute their preferred approach for the current Cufflinks integration in LESSeq, while the remainder of the pipeline remains unchanged.

### Identifying local events

In this step, LESSeq builds a splicing graph for each gene locus utilizing assembled splicing structures from the previous step (see ‘Refining gene models using RNA-Seq’). Locally diverged structures are identified in the locus graphs and defined as ‘local events’ (Fig. 1B). As illustrated in the two scenarios described in Fig. 1B, the definition of local events is conservative, so that any transcript-splicing structure from a gene locus must go through one of the paths in a local event. In the current implementation, the shortest, conservative local graphs are taken. However, the identified local events for some genes can still be very complex, and suffer from similar quantification difficulties as those in transcript-based analysis, such as unidentifiability (10). Therefore, the pipeline generates predefined and simple patterns of local events (4). In these cases, all splicing models of a gene can be grouped into one of two alternative forms at a local event. The simple event patterns, as defined in Wang et al. (2008) (4), included in LESSeq are: Skipped Exon (SE), Retained Intron (RI), Alternative 5’ Splice Site (A5SS), Alternative 3’ Splice Site (A3SS), Mutually eXclusive Exon (MXE), Alternative First Exon (AFE), Alternative Last Exon (ALE) and Tandem 3’ UTRs (T3). It should be noted that the events selected are restrictive so that downstream analyses yield robust results.

### Counting reads that are compatible with alternative forms of local events and estimating their relative expression levels

A metric to quantify isoform usage is the relative expression level of an isoform, where each isoform’s expression level is divided by the total expression from all isoforms of a given gene. As such, the relative expression level represents isoform abundance relative to other isoforms of a given gene, and the sum of all isoforms’ relative expression levels for a gene is 1.0. This metric is useful if one aims to compare alternative isoform usage independent of gene expression level changes. For the local events identified from the previous step, such a concept leads to the natural definition of relative expression levels of alternative forms for each local event (Fig. 2A). For each pre-defined simple event type, there are two fractional values representing the relative expression level of either of the two possible forms of a local event. Such metric provides a quantitative measurement of the extent of alternative splicing at each given locus, and the values can be used to examine the relationships between samples with clustering (Fig. 2B).

**Fig. 2.**
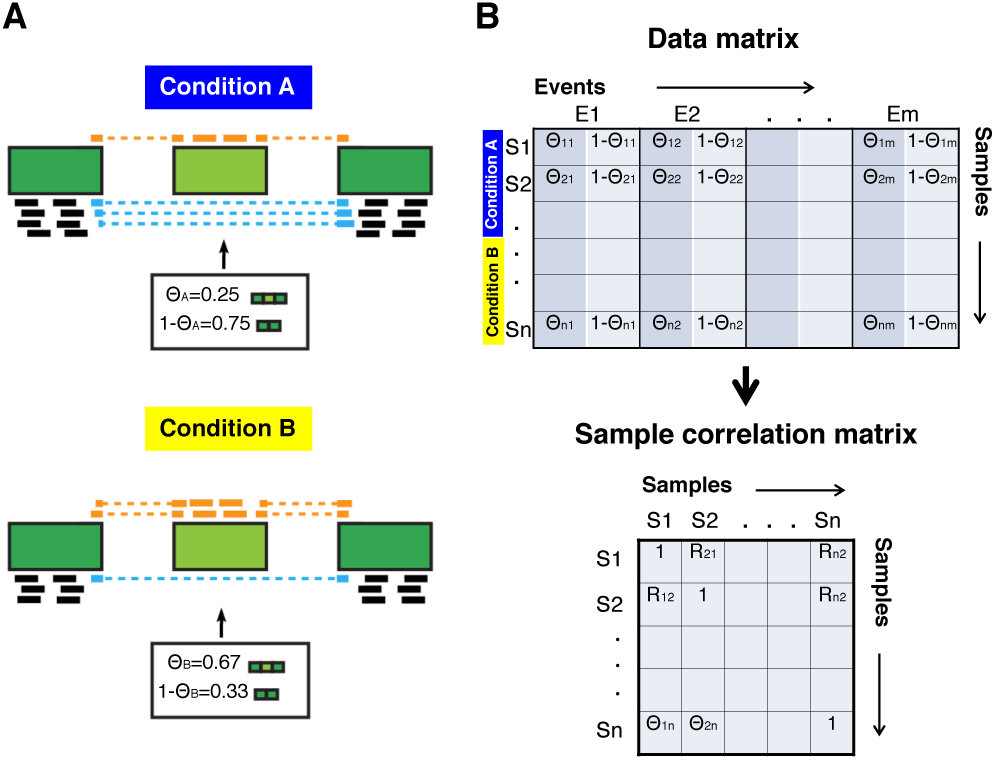
Local event expression estimation and comparative analysis. (**A**) The generative process of RNA-Seq underlying the partial probabilistic sampling-based approach to estimate relative expression levels of local events, using ‘skipped exon’ (1 − *θ*) as an example. Conditions A and B portray how the estimated parameters (i.e., relative expression levels, *θ* and 1 − *θ*) give rise to the observed RNA-Seq read distribution at local splicing events. Reads colored black are compatible with both forms of a local events, while the orange and cyan reads come from different forms of a local event. The parameter estimation procedure uses all three types of reads and builds a data matrix (panel **B** *top*) that records the compatibility information between the reads and the exons in a local event. The Maximum Likelihood Estimation (MLE) of relative expression levels is then derived, assuming uniform generation of RNA-Seq reads along each form of a local event. For differential alternative splicing testing, two hypothetical conditions are considered. Raw read counts are used in the parametric methods, while the estimated relative expression levels are used in the non-parametric method. When counting raw reads, the number of reads compatible with one form of a local event is the sum of black reads plus orange or cyan reads, with the choice of orange or cyan dependent on which form is being tested; the total number of reads in a local event is the sum of all black, orange and cyan reads. (**B**) Expression levels of local events can be used to examine the relationship between samples through clustering. The illustrated process is used to generate heatmaps in Figures 3A and 4A (*left*). A data matrix (*top*) is built from relative expression levels of all local events, and subsequently used to construct a sample correlation matrix (Spearman’s Rank-Order Correlation Coefficient, *ρ*, is used in the analyses throughout this paper). Clustering of samples is based on the sample correlation matrix. For Figure 4A, the heatmap on the *right* is generated using a similar procedure with the relative expression levels (two values for each event) replaced by the total expression levels (one values for each event).

To calculate the relative expression levels of each alternative form in a local event, LESSeq counts the number of reads that are compatible with either forms at each locus (Figure 2A) and then derives the Maximum Likelihood Estimates (MLE). MLEs are thus calculated using RNA-Seq read counts for local event annotations. LESSeq implements a similar methodology as described in Du et al. (2012) (8), which estimates relative isoform expression levels. In this methodology, RNA-Seq data is modeled as a probabilistic partial sampling process, based on a generative model of reads from alternative forms of each local event. A uniform distribution of short-reads along each isoform is assumed and an Expectation-Maximization (EM) algorithm is used to infer the MLE. The output from this step of the LESSeq pipeline is the raw number of reads associated with each form of a local event as well as the estimated relative expression levels.

### Testing differential alternative splicing events

In the final step of the pipeline, LESSeq determines the statistical significance of differential alternative splicing events between conditions. Parametric tests (two-sided Fisher exact and log-linear model for one or more replicates, respectively) are performed when very few replicates (e.g., three replicates or fewer) are available for each condition. A non-parametric test (Wilcoxon rank sum) is performed when there are many replicates per condition. In studies where many replicates are generated, the non-parametric test can supplement the parametric tests for two purposes: i) to compare the abundance of significant differential alternative splicing events and ii) to derive the most confident candidates (by taking the intersection of results from multiple tests). When only one replicate is generated for each condition, a two-way contingency table is constructed for each local event, where each cell’s value is the raw read-count compatible with one form of a local event in one condition (see Fig. 2A and Table 1).

**Table 1.**
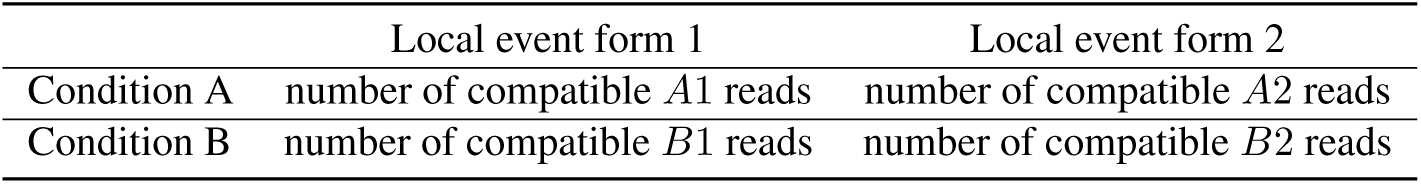
Example two-way contingency table for a two-form local event between hypothetical conditions A and B.

|  | Local event form 1 | Local event form 2 |
| --- | --- | --- |
| Condition A | number of compatible *A1* reads | number of compatible *A2* reads |
| Condition B | number of compatible *B1* reads | number of compatible *B2* reads |

When more than one replicate is available in each condition, the parametric test is based on a log-linear model fitted with a Poisson link and subsequent likelihood ratio test based on model fit (13–16). In this model, the number of raw reads compatible with one form *p* of a local event in sample *i* is denoted as *X*_*pi*_ and modeled as:

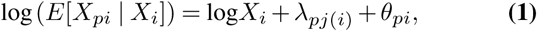

where *X*_*i*_ is the total number of raw reads mapped in the local event for sample *i, λ*_*pj*(*i*)_ is the condition-specific splicing level for condition *j* and *θ*_*pi*_ is the replicate error term. *λ*_*pj*(*i*)_ and *θ*_*pi*_ are learned by R’s built-in linear model function when fitting the data.

For each event, *p* can be either of the two forms of a local event, where two tests are performed resulting in two p-values generated for each event. The significance of differential alternative splicing for each event can be represented by either the smaller or the greater p-value (the smaller p-value was used in our analysis of RNA-Seq data). Raw read counts, compatible with one form in a simple-patterned local event, are compared between conditions and normalized for the total raw read counts in the entire locus. Normalization was performed to ensure that the effect of differential alternative splicing is tested, independent of total expression level changes.

When many replicates exist for each condition, a non-parametric test can also be applied. The non-parametric test takes as input the relative expression levels estimated (from ‘Counting reads that are compatible with alternative forms of local events and estimating their relative expression levels’), as opposed to raw read counts in the parametric tests. Using relative expression levels in each form of a local event, the method performs a Wilcoxon rank sum test on the distributions between different conditions. Here, relative expression levels for one form of a local event are represented as a vector with the length equal to the number of samples in a given condition. Two vectors representing different conditions are then compared to each other using the Wilcoxon rank sum test.

## Results

To demonstrate the usefulness of the LESSeq pipeline, we applied LESSeq to two studies of comparative RNA-Seq data: i) within a single species and ii) across multiple species.

### LESSeq identifies within-species variation of human populations from RNA-Seq data

By investigating comparative RNA-Seq experiments obtained from different conditions in a single species, LESSeq was used to identify alternative splicing signatures important to various aspects of biology. For example, RNA-Seq data could be sampled from different time points during organism development to yield insights into the genomic events driving developmental progression, from different organs to identify signatures underlying tissue differentiation or performing a comparison between healthy vs. disease-state samples to facilitate the discovery of aberrant splicing that is disease specific.

Here, we address the question of variation between individuals and populations. To accomplish this, LESSseq was used to study the differences in alternative splicing found within a data set generated by the Geuvadis Consortium (17). Mapped messenger RNA (mRNA) reads for RNA-Seq data in Lymphoblastoid Cell Lines (LCL) of two human populations (CEU and YRI, with 91 and 89 samples, respectively) were downloaded (http://www.ebi.ac.uk/arrayexpress/files/E-GEUV-1/processed/). Gene annotation refinement using RNA-Seq reads and the EnsEMBL (V67) (18) human annotation led to 2,948 local simple-patterned events being identified. Each event was required to have no less than 80nt-long exons and 50nt-long introns. By using the relative expression levels estimated for all local events, samples were clustered and revealed that individuals do not segregate by population with regard to alternative splicing in these local events (Fig. 3A). Statistical tests for alternative splicing changes yield between 8% to 10% differential events between the two populations, with 174 events detected by both parametric and non-parametric methods (Benjamini-Hochberg corrected p-value, or BHP cutoff at 0.05, Figs. 3B and 3C).

**Fig. 3.**
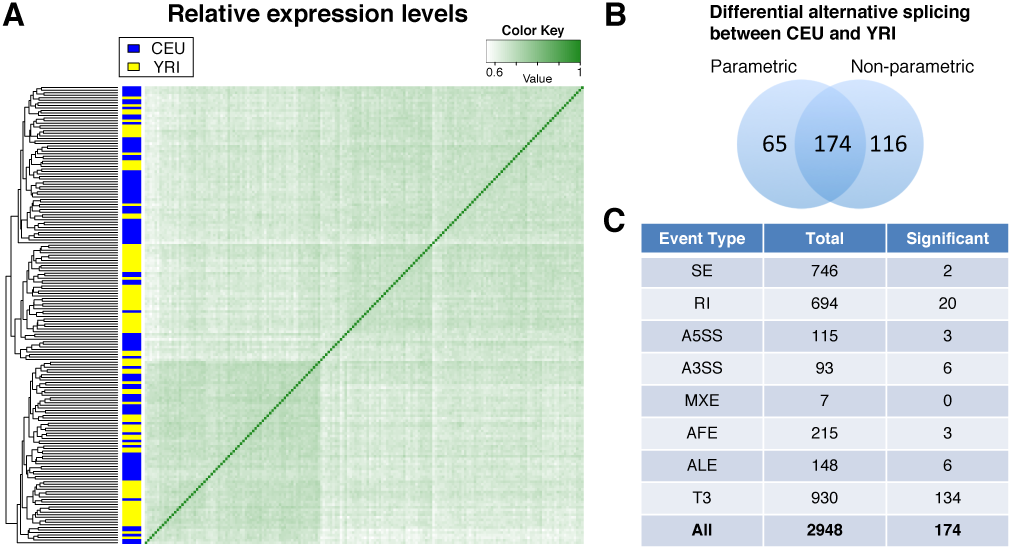
Analysis of variation between human individuals. (**A**) Complete linkage hierarchical clustering of 91 CEU and 89 YRI samples from Lappalainen et al. (2013) (17), using the sample correlation matrix calculated from relative expression levels of 2,948 local events. Distance between samples was measured as 1-*ρ*. Individuals were shown not to cluster by population identity as demonstrated by original authors (17). (**B**) Overlap of the significant differential alternative splicing events (BHP cutoff 0.05) found by the log-linear model method (‘parametric’) and the Wilcoxon rank sum test (‘non-parametric’). (**C**) Number of events identified (‘Total’) and those found to be significant by both methods (‘Significant’), broken down by event type: SE - skipped exon, RI - retained intron, A5SS - alternative 5’ splice site, A3SS - alternative 3’ splice site, MXE - mutually exclusive exon, AFE - alternative first exon, ALE - alternative last exon, T3- tandem 3’ UTRs and All - the sum of all event types.

In the original paper, Lappalainen et al. (2013) (17) demonstrated that populations can be clustered by genotype, but not by exon-level expression. Exon-level expression is the combined product of gene expression level and alternative splicing. It is not informative to assess these two aspects of gene regulation separately. Using local event relative expression levels, we were able to examine quantitatively the variations of alternative splicing alone at the identified local events. Our clustering results showed that individuals do not cluster by population in this regard (Fig. 3A). The original research paper also attempted to identify population-differential alternative splicing events. Using GENCODE (19) annotations, transcript-based (Flux-capacitor (20) and Wilcoxon rank sum test for transcript quantification and significance testing, respectively) and exon-based (DEXSeq (21)) analyses were conducted, where approximately 20% and 45% significant genes were found, respectively. However, these analyses were based on reference gene annotation, and the transcript-based and exon-based methods suffer from a number of the issues discussed in the introduction and discussion sections.

When we used LESSeq to find population-differential alternative splicing events, a much smaller fraction of confident events was found. This decrease highlights the difference between local event-based and the original author’s calculations.

### LESSeq provides a statistical framework to detect quantitatively differential alternative splicing patterns across species

Comparative RNA-Seq experiments across species improves our understanding of organismal evolution in terms of transcriptome variation and aids in the identification of candidate genes that may underlie phenotypic evolution (i.e., differentially expressed and/or spliced genes). LESSeq’s approach to differential alternative splicing events was shown to achieve robust results when studying cross-species alternative splicing. In such context, transcript-based approaches are affected by non-uniform gene annotations, which leads to increased false positive detection in differential splicing. In addition, analyses that focus on reads that span splice junctions alone do not contain as much information from RNA-Seq data as LESSeq’s local event-based approach.

To study the variation of alternative splicing between human and other primates, we utilized messenger RNA (mRNA) data, observed by RNA-Seq, containing six profiled organs (brain, cerebellum, heart, kidney, liver and testis) in ten species (22). SRA files (GEO:GSE30352) were downloaded for all six organs of human and rhesus macaque (two replicates were obtained for each organ in each species). Reads were then aligned using TopHat (23). UCSC’s LiftOver tool (https://genome.ucsc.edu/cgi-bin/hgLiftOver) (24) was used to match orthologous exon coordinates between one-to-one orthologous genes (EnsEMBL’s V67 annotation was used again for both species), which yielded 1,683 orthologous ‘skipped exon’ local events at a reciprocal mapping rate of 0.9. Based on the relative expression levels calculated by LESSeq at these events, as well as total expression levels (measured by total read-counts), hierarchical clustering results show that alternative splicing patterns cluster by species whereas total expression levels cluster by tissues (Fig. 4A). Lineage-differential alternative splicing events were also identified in each tissue (Fig. 4B).

**Fig. 4.**
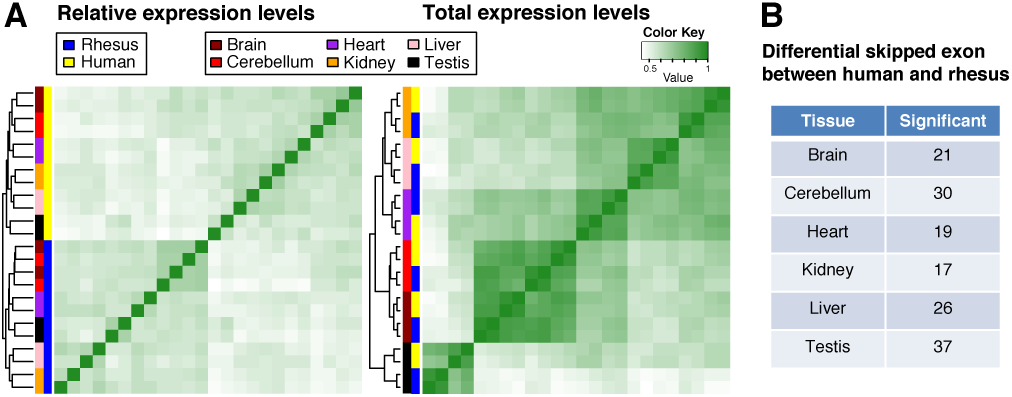
Analysis of variation between human and rhesus macaque. (**A**) Complete linkage hierarchical clustering of all 24 samples (two species, six tissues, two replicates per species-tissue combination) using the sample correlation matrix calculated from expression levels at 1,683 ‘skipped exon’ events. Distance between samples was measured as 1-*ρ*. Relative expression levels cluster by species identity (*left*), and total expression levels cluster by tissue identify (*right*). (**B**) Number of significant, differential ‘skipped exon’ events (Benjamini-Hochberg corrected p-value cutoff of 0.05) detected in each tissue between the two species, using log-linear model method (non-parametric method was not applicable due to the small number of replicates). Our results support the original authors (22) findings that a strong selection pressure exists for maintaining total gene expression levels of organs as well as recent studies (25, 26) that found faster cross-species changes in alternative splicing compared to total expression level.

Brawand et al. (2011) (22) originally revealed a strong selection pressure exists for maintaining total gene expression levels of organs. Our study confirms this by showing that the total expression levels at local events cluster by tissue identity. Additionally, we find that the relative expression levels cluster by species. This observation agrees with two recent studies that found faster cross-species changes in alternative splicing compared to total expression level (25, 26). In addition to the global observation of alternative splicing evolution, our method provides a statistical framework that can be used to detect quantitatively differential alternative splicing patterns across species – by identifying these events, we provide candidates for generating lineage-specific phenotypes.

## Discussion

In this paper, we described a pipeline (LESSeq) for the analysis of comparative alternative splicing events from RNA-seq data. LESSeq provides robust differential alternative splicing detection and uncovers localized candidate regions in genes that exhibit differential alternative splicing. By pinpointing specific loci of interest, LESSeq can ease the design of downstream mechanistic studies (e.g., mini-gene assay). The LESSeq pipeline also allows unambiguous design of PCR primers and microarray probes for large-scale applications (e.g., healthy vs. disease state biomarkers).

LESSeq employs a local event-based analysis strategy that is more robust to transcript annotation as well as quantification errors compared to transcript-based methods. Other methods have also been built around similar ‘localized’ ideas, such as DEXSeq (21) — a method that tests differential exon usage. However, compared to LESSeq, DEXSeq loses information on the local connectivity of exons and does not define the differential usage of exons as a result of different modes (e.g., skipped exon in Fig. 1B). DEXSeq also does not provide the relative expression level estimation, which is a useful metric to assess the degree of alternative splicing. Most importantly, the estimation of each exon’s expression level in DEXSeq may be confounded by other alternative splicing events found within the same gene. In comparison, the estimation of each local event in LESSeq is not affected by other events due to the strict definition of included events in the analysis (Fig. 1B). DiffSplice (27) is a method that both quantifies and tests for differential local events, or ‘alternative splicing modules’. Compared to LESSeq, DiffSplice does not filter out complex local events. The results of analyzing these complex events could be very unreliable, as some of them are computationally unidentifiable using short-read data (10). Furthermore, DiffSplice uses a permutation-based test that is not applicable when there are fewer than three replicates per condition. In such situations, LESSeq provides parametric tests that can handle very few replicates. When many replicates are available, LESSeq’s parametric and non-parametric tests can both be utilized, which provides the most confident candidates identified by the intersection of both methods (28). Mixture-of-ISOforms (MISO) (9) is another tool that can analyze local variable regions of gene structures. However, MISO requires user-supplied or curated alternative splicing annotations and does not provide the option for a user to generate variable region annotations (i.e., integrate different gene transcript models). Furthermore, it does not provide a statistical framework for differential alternative splicing testing when multiple replicates are available – a user must pool reads from replicates of one condition together, in which one discards the information on sample variations and is not an ideal statistical practice.

## Conclusions

In this paper, we presented LESSeq, a computational pipeline that infers different RNA splicing events from RNA-Seq data. LESSeq takes advantage of information found in the RNA-Seq read distribution at local splicing events (as defined in in Wang et al., 2008 (4)) to determine unambiguous alternative splicing. Unlike other transcript-based methods, LESSeq is robust to transcription annotation and allows for the integration of different gene transcript models. We demonstrated the usefulness of LESSeq by applying the pipeline to two RNA-Seq data sets containing: i) multiple human individuals (17) and ii) multiple species and organs (22). In the first data set, LESSeq identified increased variation within individuals of the same population, revealing that exon-level expression contains more information than genotype alone (used by the original authors — Lappalainen et al., 2013 (17)). Finally, applying LESSeq to RNA-Seq data from six organs across ten species supported the original authors (Brawand et al., 2011 (22)) findings that a strong selection pressure exists for maintaining gene expression levels of organs. Based on our results and the popularity of RNA-Seq, we see LESSeq becoming a useful tool for the transcriptomic community.

## Supporting information

To illustrate the advantage of using LESSeq, we simulated single-end read distributions for mammalian genes under various cellular conditions. For each example gene, 100 simulations at four sequencing depths (100, 1000, 10000 and 100000 reads) were generated, where the simulated single-end reads were 75 bp long and distributed uniformly across each transcript.

---

**Fig. S1.**
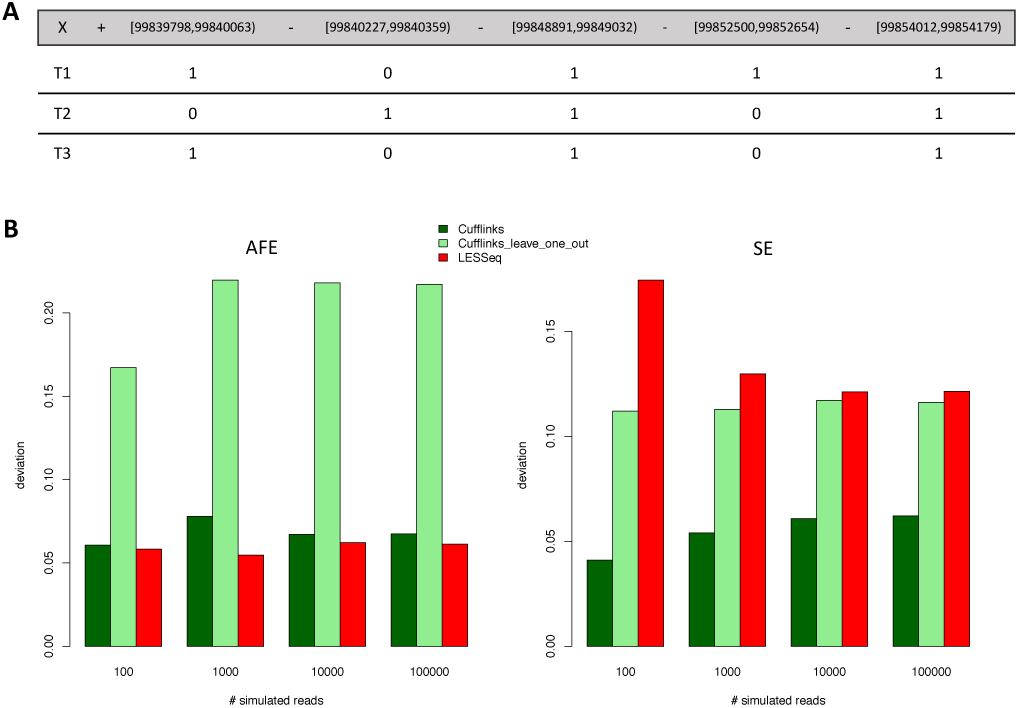
Simulation on an unidentifiable gene model. (**A**) Splicing graph for gene G1. Top row (grey background) indicates the genomic coordinates of G1 - including chromosome, strand, as well as the start and end of G1’s exons. Each exon is zero-index, where the left and right coordinates are inclusive and exclusive, respectively. T1 to T4 are the Transcript (T) names. For each transcript, ‘0’ and ‘1’ denote whether an exonic region is excluded or included within the transcript, respectively. This gene is identifiable using single-end reads because the binary matrix representing the transcript structures is full rank. (**B**) The average quantification errors at local splicing events by three methods: i) ‘Cufflinks’ - Cufflinks (11) observes all four transcripts and derives the local event quantification by summing transcript-level expression, ii) ‘Cufflinks_leave_one_out’ – Cufflinks is run on the first two transcripts (T1 and T1) and derives the local event quantification by summing transcript-level expressions, iii) ‘LESSeq’ - LESSeq is used to identify and quantify local events. The relative expression level of each transcript is set to be the same (1/3). For each of the two local events in this gene, the absolute difference between the average of 100 simulations and simulation setting was calculated and plotted on Y-axis. ‘Cufflinks’ performs best because all the true transcript structures are provided and this method has the advantage of taking all reads into account when deriving the local event quantification. However, when the correct transcript structures are not given, which occurs quite frequently (6), LESSeq performs better than ‘Cufflinks_leave_one_out’ for the Alternative First Exon (AFE) event and is comparable for the Skipped Exon (SE) event. The ‘Cufflinks_leave_one_out’ mode performs unsatisfactorily at local events because G1 is identifiable (unlike G2 in Supplemental Fig. S2), while the transcript-level quantification is very poor (Supplemental Table S1).

**Table S1.** Average transcript-level estimations for G1.

| Method | Number of reads | T1 | T2 | T3 |
| --- | --- | --- | --- | --- |
| Cufflinks | 100 | 0.31885 | 0.42154 | 0.25961 |
|  | 1000 | 0.27413 | 0.41721 | 0.30866 |
|  | 10000 | 0.26864 | 0.40253 | 0.32883 |
|  | 100000 | 0.27126 | 0.40082 | 0.32792 |
| Cufflinks_leave_one_out | 100 | 0.47196 | 0.52804 |  |
|  | 1000 | 0.44097 | 0.55903 |  |
|  | 10000 | 0.44662 | 0.55338 |  |
|  | 100000 | 0.44955 | 0.55045 |  |

**Table S2.** Average transcript-level estimations for G2.

| Method | Number of reads | T1 | T2 | T3 | T4 |
| --- | --- | --- | --- | --- | --- |
| Cufflinks | 100 | 0.00000 | 0.29994 | 0.39021 | 0.30985 |
|  | 1000 | 0.00000 | 0.00000 | 0.51325 | 0.48675 |
|  | 10000 | 0.00000 | 0.03138 | 0.49556 | 0.47306 |
|  | 100000 | 0.00000 | 0.00000 | 0.51409 | 0.48591 |
| Cufflinks_leave_one_out | 100 | 0.29283 | 0.62843 | 0.07874 |  |
|  | 1000 | 0.42122 | 0.55674 | 0.02204 |  |
|  | 10000 | 0.41129 | 0.57045 | 0.01826 |  |
|  | 100000 | 0.41070 | 0.56651 | 0.02279 |  |

**Fig. S2.**
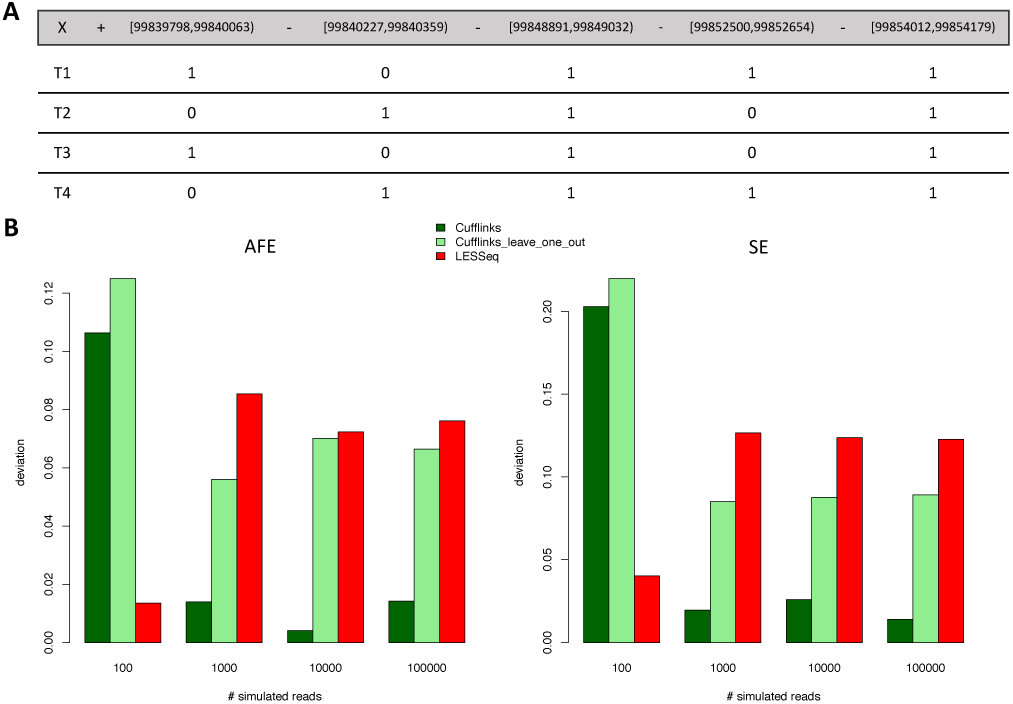
Simulation on an unidentifiable gene model. (**A**) Splicing graph for gene G2 in the same format as Supplemental Fig. S1. G2 is unidentifiable using single-end reads, because the binary matrix representing the transcript structures is not full rank. (**B**) The average quantification errors at local events by three methods: i) ‘Cufflinks’ - Cufflinks (11) observes all four transcripts and derives the local event quantification by summing transcript-level expression, ii) ‘Cufflinks_leave_one_out’ – Cufflinks is run on the first two transcripts (T1 and T1) and derivES the local event quantification by summing transcript-level expressions, iii) ‘LESSeq’ - LESSeq is used to identify and quantify local events. The relative expression level of each transcript is set to be the same (1/4). For each of the two local events in G2, the absolute difference between the average of the 100 simulations and the simulation setting was calculated and plotted on Y-axis. In this example, LESSeq performs slightly worse than both ‘Cufflinks’ and ‘Cufflinks_leave_one_out’. When Cufflinks is provided all ‘true’ transcript structures (‘Cufflinks’), this method has the advantage of taking all reads into account when calculating transcript-level expression (though the transcript-level quantification is quite poor – see Supplemental Table S2). ‘Cufflinks_leave_one_out’ (T4 is left out in the annotation) is shown to perform decently because T3 is always assigned a small expression level (Supplemental Table S2). Therefore, the local splicing event quantification is not dramatically affected by inaccurate transcript annotation (due to the similarity between T1 and T3) even though the transcript-level quantification is poor.

## Bibliography

1. Nilsen, T. & Graveley, B. Expansion of the eukaryotic proteome by alternative splicing. Nature 463, 457–463 (2010).

2. Lu, Z., Jiang, P. & Xing, Y. Genetic variation of pre-mRNA alternative splicing in human populations. Wiley Interdiscip Rev RNA 3, 581–592 (2012).

3. Wang, Z., Gerstein, M. & Snyder, M. RNA-Seq: a revolutionary tool for transcriptomics. Nat Rev Genet 10, 57–63 (2009).

4. Wang, E. et al. Alternative isoform regulation in human tissue transcriptomes. Nature 456, 470–476 (2008).

5. Pickrell, J., Pai, A., Gilad, Y. & Pritchard, J. Noisy splicing drives mRNA isoform diversity in human cells. PLoS Genet 6, e1001236 (2010).

6. Steijger, T. et al. Assessment of transcript reconstruction methods for RNA-seq. Nat Methods 10, 1177–1184 (2013).

7. Pachter, L. Models for transcript quantification from RNA-Seq. arXiv 1104.3889v2 (2011).

8. Du, J. et al. IQSeq: integrated isoform quantification analysis based on next-generation sequencing. PLoS One 7, e29175 (2012).

9. Katz, Y., Wang, E., Airoldi, E. & Burge, C. Analysis and design of RNA sequencing experiments for identifying isoform regulation. Nat Methods 7, 1009–1015 (2010).

10. Hiller, D., Jiang, H., Xu, W. & Wong, W. Identifiability of isoform deconvolution from junction arrays and RNA-Seq. Bioinformatics 25, 3056–3059 (2009).

11. Trapnell, C. et al. Transcript assembly and quantification by RNA-Seq reveals unannotated transcripts and isoform switching during cell differentiation. Nat Biotechnol 28, 511–515 (2010).

12. Garber, M., Grabherr, M., Guttman, M. & Trapnell, C. Computational methods for transcriptome annotation and quantification using RNA-seq. Nat Methods 8, 469–477 (2011).

13. Ayoub, A. et al. Transcriptional programs in transient embryonic zones of the cerebral cortex defined by high-resolution mRNA sequencing. Proc Natl Acad Sci U S A 108, 14950–14955 (2011).

14. Bullard, J., Purdom, E., Hansen, K. & Dudoit, S. Evaluation of statistical methods for normalization and differential expression in mRNA-Seq experiments. BMC Bioinformatics 11, 94 (2010).

15. Cotney, J. et al. Chromatin state signatures associated with tissue-specific gene expression and enhancer activity in the embryonic limb. Genome Res 22, 1069–1080 (2012).

16. Cotney, J. et al. The evolution of lineage-specific regulatory activities in the human embryonic limb. Cell 154, 185–196 (2013).

17. Lappalainen, T. et al. Transcriptome and genome sequencing uncovers functional variation in humans. Nature 501, 506–511 (2013).

18. Flicek, P. et al. Ensembl 2014. Nucleic Acids Res 42, D749–D755 (2014).

19. Harrow, J. et al. GENCODE: the reference human genome annotation for The ENCODE Project. Genome Res 22, 1760–1774 (2012).

20. Montgomery, S. et al. Transcriptome genetics using second generation sequencing in a Caucasian population. Nature 464, 773–777 (2010).

21. Anders, S., Reyes, A. & Huber, W. Detecting differential usage of exons from RNA-seq data. Genome Res 22, 2008–2017 (2012).

22. Brawand, D. et al. The evolution of gene expression levels in mammalian organs. Nature 478, 343–348 (2011).

23. Trapnell, C., Pachter, L. & Salzberg S. TopHat: discovering splice junctions with RNA-Seq. Bioinformatics 25, 1105–1111 (2009).

24. Hinrichs, A. et al. The UCSC Genome Browser Database: update 2006. Nucleic Acids Res 34, D590–D598 (2006).

25. Barbosa-Morais, N. et al. The evolutionary landscape of alternative splicing in vertebrate species. Science 338, 1587–1593 (2012).

26. Merkin, J., Russell, C., Chen, P. & Burge C. Evolutionary dynamics of gene and isoform regulation in Mammalian tissues. Science 338, 1593–1599 (2012).

27. Hu, Y. et al. DiffSplice: the genome-wide detection of differential splicing events with RNA-seq. Nucleic Acids Res 41, e39 (2013).

28. Soneson, C. & Delorenzi, M. A comparison of methods for differential expression analysis of RNA-seq data. BMC Bioinformatics 14, 91 (2013).

